# Predicting microbiomes through a deep latent space

**DOI:** 10.1101/2020.04.27.063974

**Authors:** Beatriz García-Jiménez, Jorge Muñoz, Sara Cabello, Joaquín Medina, Mark D. Wilkinson

**Affiliations:** Centro de Biotecnología y Genómica de Plantas (CBGP, UPM-INIA), Universidad Politécnica de Madrid (UPM) – Instituto Nacional de Investigación y Tecnología Agraria y Alimentaria (INIA), Campus de Montegancedo-UPM, 28223, Pozuelo de Alarcón (Madrid), Spain; Serendeepia Research, 28905 Getafe (Madrid), Spain

## Abstract

**Motivation:** Microbial communities influence their environment by modifying the availability of compounds such as nutrients or chemical elicitors. Knowing the microbial composition of a site is therefore relevant to improving productivity or health. However, sequencing facilities are not always available, or may be prohibitively expensive in some cases. Thus, it would be desirable to computationally predict the microbial composition from more accessible, easily-measured features.

**Results:** Integrating Deep Learning techniques with microbiome data, we propose an artificial neural network architecture based on heterogeneous autoencoders to condense the long vector of microbial abundance values into a deep latent space representation. Then, we design a model to predict the deep latent space and, consequently, to predict the complete microbial composition using environmental features as input. The performance of our system is examined using the rhizosphere microbiome of Maize. We reconstruct the microbial composition (717 taxa) from the deep latent space (10 values) with high fidelity (¿0.9 Pearson correlation). We then successfully predict microbial composition from environmental variables such as plant age, temperature or precipitation (0.73 Pearson correlation, 0.42 Bray-Curtis). We extend this to predict microbiome composition under hypothetical scenarios, such as future climate change conditions. Finally, via transfer learning, we predict microbial composition in a distinct scenario with only a hundred sequences, and distinct environmental features. We propose that our deep latent space may assist microbiome-engineering strategies when technical or financial resources are limited, through predicting current or future microbiome compositions.

**Availability:** Software, results, and data are available at https://github.com/jorgemf/DeepLatentMicrobiome

## 1 Introduction

### The microbiome

Microbes are everywhere, in human, animals, plants and the environment (soil, water, air), executing numerous biological functions whose absence would dramatically reduce the quality and quantity of life on earth (Gilbert and Neufeld, 2014). Study of microbial communities has increased in recent years due to advances in high-throughput technologies that now allow identification of microbes in a community by sequencing rather than culturing (Lloyd-Price *et al.*, 2017; Thompson *et al.*, 2017). Microbial community functions include collaborating in carbon and nitrogen cycles, to provide nutrients to animal and plant cells by breaking complex molecules into smaller compounds, training and triggering the immune system to fight against pathogens, etc. Those microbiome functions entail applications in health and medicine, climate change, sustainable agriculture, environment and biofuels.

Most microbiome analyses to date have focused on observational or descriptive approaches; i.e. to identify the microbes living in a community and to establish correlations between those experimental findings and some phenotypic feature, such as abundance or deficiency of a particular strain in a disease’s subject group. For example, comparing human gut in health vs gastrointestinal disease subjects (Kotloff *et al.*, 2013) or the soil microbiome before and after a large environmental challenge (Uritskiy *et al.*, 2019). Some microbiome studies go one step further, following a predictive approach. These translational approaches should allow us to design solutions based on microbiome modulation to address problems in human and plant health, and take microbial composition as a predictor of a particular phenotypic feature, using linear regression and machine learning (ML) techniques (Caporaso *et al.*, 2011; Zhou and Gallins, 2019).

### ML in bioinformatics and microbiome research

Machine Learning (ML) has been utilized in bioinformatics analyses for many years. Areas of application include sequence analysis, structure and function prediction, and interaction prediction (Inza *et al.*, 2010).

Studies focused on the microbiome and metagenomics, where ML approaches are applied, have recently grown in number (Bokulich *et al.*, 2018; Sakowski *et al.*, 2019). We distinguish two major types of study. First, work on single cases, where a unique dataset or problem is addressed, for example, microbial composition being used to predict productivity in soil (Chang *et al.*, 2017), contaminants and geochemical features in wells (Smith *et al.*, 2015), presence/absence of disease due to changes in abundances of microbes over time (Bogart *et al.*, 2019), or biomarkers of cancer (and the type of cancer) from the human blood microbiome (Poore *et al.*, 2020). Second, more general studies are emerging, where multiple datasets of different origin are addressed together, applying the same prediction procedure or tool. For example, MetAML (Pasolli *et al.*, 2016), a computational tool for metagenomics-based prediction tasks and for microbiome-phenotype associations; q2-sample-classifier (Bokulich *et al.*, 2018), a plugin for QIIME 2 for supervised classification; and Microbiome Learning Repo (ML Repo) (Vangay *et al.*, 2019) which provides a benchmark of 33 curated classification and regression tasks from 15 published human microbiome datasets.

### Deep Learning in bioinformatics and microbiome/metagenomics

Recently, there has been an emergence of Deep Learning (DL) approaches (Lecun *et al.*, 2015) to biological challenges (Min *et al.*, 2017; Ching *et al.*, 2018). The areas of application span those of the classical ML approaches listed above. DL has generated performance improvements in areas where the input data are images or sequences (i.e. DL propitious data formats), such as diagnosis by biomedical image analysis and prediction based on sequence analysis (Li *et al.*, 2019). DL also contributes, in the bioinformatics domain, to improvements in data representation or automatic feature extraction, for example retrieving features from Electronic Health Records to calculate a patient’s risk of disease (Miotto *et al.*, 2016).

With respect to microbiome studies, DL has not yet been as widely applied compared to other bioinformatics problems, and thus examples are limited: phenotype prediction (food allergy) from bacterial composition with LSTM (Metwally *et al.*, 2019), human age prediction (Galkin *et al.*, 2018), identification of body-site and prediction of Crohn’s disease with a k-mer representation (Asgari *et al.*, 2018), or identification of microbiome biomarkers using graph embedding (Zhu *et al.*, 2019). Generally speaking, phenotype prediction from metagenomics data is the most common task solved by M/DL in the microbiome space (LaPierre *et al.*, 2019).

### Our proposal

Pursuing the promising translational and predictive approach, in this work we propose an ambitious goal beyond predicting phenotypic features from microbial composition. Rather, we attempt to achieve the opposite: to predict the microbial composition based on a few phenotypic and/or environmental features. Our approach would be useful, for example, in cases where one must make a decision whose outcome depends on microbial composition, but where sequencing facilities are not available, the sequencing cost is prohibitive, or where the only available data are phenotypic/environmental features. For example, a farmer in an emerging economy, with no access to sequencing facilities (nor finances to engage them), might want to make strategic decisions about what crop to cultivate on their land, or which fertilizer to use in what amount – a decision highly dependent on that soil’s microbiome, which determines the nutritional capabilities of that soil.

Few previous studies have tried to predict microbial composition from environmental features, and those have mainly focused on a few tens of taxa, corresponding to the highest taxonomic levels (Larsen *et al.*, 2012; Ladau *et al.*, 2018; Oh and Zhang, 2020).

### Our approach and contributions

The primary contributions of this work are: a) the development of a novel autoencoder model that merges knowledge from both microbial composition and environmental/mapping features, into a deep latent space; b) an exhaustive selection and evaluation of autoencoder reconstructions of the whole microbial composition (hundreds of taxa) from that latent space; and c) prediction of microbiome composition starting from only environmental features. All of these contributions are supported by real data from a soil microbiome study, in particular, the maize rhizosphere microbiome. Maize is an important food crop, in a world reaching an estimated 10 billion people by 2050. This will require a doubling of food production on scarce agriculture soil, with increasingly limited water, and avoiding the expensive and destructive application of fertilizers (Hunter *et al.*, 2017). As such, agrifood research – the focus of this work – is of great interest and socioeconomic importance.

While the initial application of our system is aimed at predicting the microbiome based on a limited number of environmental features, we believe that the system could also be applied to the prediction of the microbiome for novel or hypothetical ecosystems. This would help us prepare for the consequences of environmental influences such as climate change, or toxic spills. In addition, our system can be applied to predict the microbiome of novel datasets, with a limited number of samples or observations, via transfer learning.

## 2 Materials and Methods

### 2.1 Dataset

We selected the soil microbiome of an agronomically important plant (Zea mays L. subsp. mays), using a dataset taken from the Walters *et al.* (2018) study. The authors examined the influence of the soil microbiome on the traits of maize to clarify whether the surrounding microbial communities could be used as a breeding trait. This 16S rRNA dataset includes multiple cropping fields, and includes 27 different maize varieties.

The mapping data includes 72 features such as sequences and geo-localization. Additional metadata about environmental factors such as temperature and precipitation was kindly provided to us by the authors as it was not included in their online repository. From these, we selected 5 environmental features based on prior knowledge of factors likely to influence the plant-associated soil microbiome, excluding geographical location features to ensure the predictor was location-independent. The selected factors were: temperature, precipitation (accumulation 3 previous days), plant age, maize line and maize variety.

The initial number of OTUs from the study was 29,689, but the authors (Walters *et al.*, 2018) reduced this to 717 OTUs, corresponding to the OTUs shared in at least 80 percent of the samples, and samples with at least 10000 reads. We further filtered-out approximately 100 bulk soil samples, thus our final dataset consists of 4724 samples × 717 OTUs × 5 environmental features. Samples were split into training:testing (90:10%) sets, and within the training set there was 5-fold Cross-validation (CV).

In addition, we took a smaller dataset from the Maarastawi *et al.* (2018) study, focused on Italian and Philippine maize rhizosphere microbiomes. This small dataset was selected to become our transfer learning proof-of-concept. The original dataset has 322 samples and 1943 bacterial OTUs. After filtering out samples from bulk soil and missing values in meta-data, 123 samples remained. Regarding OTUs, due to lack of standardization in microbiome taxa annotation, it was infeasible to map those 1943 OTUs to the 717 OTUs identified in the Walters *et al.* (2018) dataset. There was limited overlap at low taxonomic levels (26/45 Class, 36/83 Order, 70/144 Family, 84/222 Genus and 0/717 Species level). The final dataset therefore included only 15 of 16 OTUs, shared at the Phylum level, that could be used for transfer learning studies. The mapping features in this case correspond primarily to chemical soil properties (pH, Nitrogen and Carbon concentrations, clay fraction, soil type, and water holding capacity). In this small dataset, samples were split into training:testing (70:30%) sets.

### 2.2 Model architectures

Our DL models follow an Auto-Encoder (AE) architecture. As such, we first design an encoder model that transforms the input features (in this case, microbial abundances and/or environmental features) to a latent space. Subsequently, we design a decoder model that predicts the output (in this case, the microbial composition) from the latent space (see Figure 1). The latent space is an encoded representation of the input features, that (generally) reduces their dimensionality; for example, in our case, from ¿700 OTUs plus the environmental variables down to ten values. Those values, in the latent space, are non-linear combinations of the input features.

**Figure 1:**
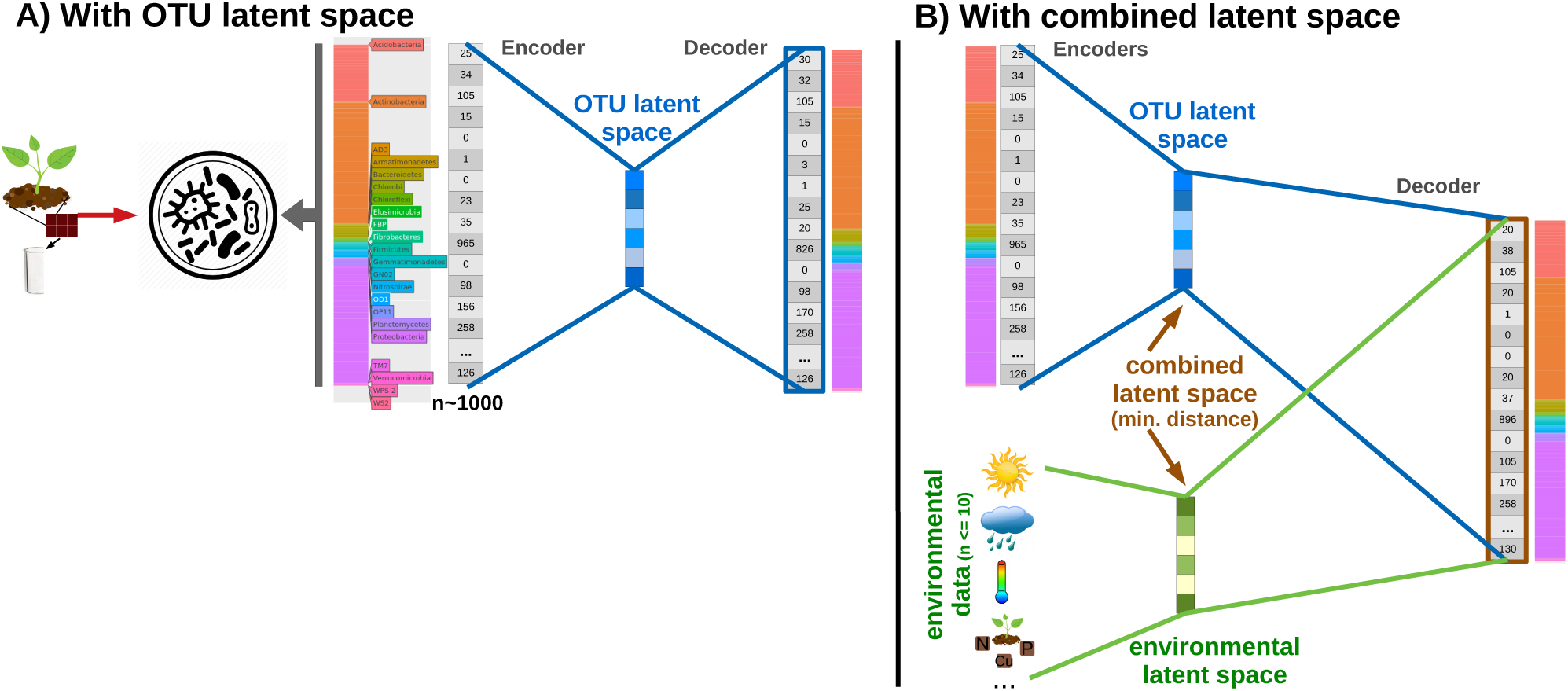
Schema of autoencoder (AE) architectures. A) AE architecture with an OTU latent space. B) AE architecture with a combined latent space (brown), which minimizes the distance between OTU (blue) and environmental (green) latent spaces during model training.

We defined two encoders:

- OTU composition → latent space
- environmental variables → latent space and one decoder model:
- latent space → OTU composition

With those modules, we then designed two different AE architectures: In one AE, the latent space depends only on the OTU composition encoder (Figure 1A). In the second AE, we have two different encoders, one for the OTU composition and another one for the environmental features. Both encoders share the same decoder. During training, the latent space of both encoders is force to be similar, this is, the difference of both latent spaces is minimized to be as close to 0 as possible (Figure 1B). Both AE architectures allow prediction of the OTU composition using only environmental features – the final goal of this study.

### 2.3 Normalization

We selected two normalization approaches suitable to 16S microbiome absolute abundance data (Weiss *et al.*, 2017), and satisfying the constraints of our DL model based on an autoencoder, which requires inverse transformations to reconstruct the original input.

The first is Total-Sum normalization (TSS), i.e. relative abundances. This approach is commonly used in DL (Oh and Zhang, 2020), although it brings with several disadvantages, in that it does not remove compositionality (Aitchison, 1982). As such, several standard analyses cannot be applied without bias, such as comparative analysis between groups; however, these problems are distinct from how we use this approach in our study.

The second approach is TSS followed by Centered Log Ratio (CLR). A logarithmic transformation is recommended to remove the compositionality of microbiome data (Zhou and Gallins, 2019). Because of the incompatibility of OTU table microbiome data sparsity (i.e. a high number of zeros) and logarithmic analyses, a small positive value is added to all abundances (in our case, pseudo-count = 1*e*^−6^). This normalization approach provides interesting advantages in our DL architecture. Although CLR can return negative values, the autoencoder accepts these as input, without substituting them with zeros (which, in other approaches, has the consequence of confounding them with the original structural zeros). Our AE architecture requires an inverse function, and this is provided by CLR in the form of Softmax (Pawlowsky-Glahn *et al.*, 2015). Moreover, Softmax is a standard activation function in DL architecture used to provide values similar to a probability distribution (all values add 1), simplifying the transformation of our output.

### 2.4 Loss functions

We have used different loss functions in our experiments to guide the training.

- *Mean Squared Error (MSE)*. The MSE is commonly used in AE as one the basic reconstruction error functions. It does not take into account information about the environmental features and it only tries to minimize the difference between the input and the output, giving more relevance to the bigger errors.
- *Crossentropy*. In our problem, it is more important to predict the proportions of each OTU rather the concrete values. So, if we use relative frequencies for the OTUs we can use statistical distances as the Crossentropy. It has been used extensively for DL in the last years. For some experiments, we decided that the output of the AE was the relative frequency of the OTUs, which allowed us to use Crossentropy directly. For the AEs where we used the CLR for the normalization we apply the Softmax transformation to the output to get the relative frequencies.
- *Bray-Curtis dissimilarity*. Since we are analysing microbiome data, we also consider community ecology approaches as loss function, selecting a commonly used microbiome beta-diversity (i.e. between-samples distance) metric – the Bray-Curtis dissimilarity. Other metrics were considered to be the loss function but they are not differentiable.

### 2.5 Evaluation metrics

We use several kinds of metrics to assess the performance of our prediction model, comparing the predicted vs actual microbial composition. First, the most common metrics used in regression learning tasks to assess reconstruction error in autoencoders are the Mean Absolute Error (MAE), Mean Squared Error (MSE) and Mean Absolute Percentage Error (MAPE). However, these are difficult to interpret (beyond ‘the lowest’ and ‘the best’) and in this case would be difficult to compare because we apply different normalization approaches, confounding these metrics because they are scale-dependent. As such, we compute additional scale-independent metrics, such as Pearson correlation, that appear also in other bio-autoencoder systems (Manica *et al.*, 2019). An additional metric selected was Bray-Curtis dissimilarity, described above in Loss Functions section.

To sort the best-predicted OTUs, we use the Root Relative Squared Error (RRSE) – a scale-independent error metric that could be applied to individual OTUs, rather than samples, and is suitable for comparing variables with large differences in values, as seen in the relative abundances of different taxa.

### 2.6 Parameterization: Model training and selection

The hyperparameters of the DL model were selected according to the evaluation metrics over a 5-fold CV within the training data set. Apart from the normalization and loss described above, the other parameters whose values were tuned were: the latent space size (10, 50, 100), number and size of the hidden layers ([512, 256], [256], none), the activation function of the encoder and decoder (tanh, relu, sigmoid) and of the latent layer (tanh, sigmoid), the learning rate (0.01, 0.001) and batch size (64, 128). Additional details are in the Jupyter notebook available on GitHub.

The experiments for hyper-parameter selection were run on a Linux server with two Nvidia 1080Ti GPUs with 11 GB of memory each, a Intel CPU 6900 with 16 cores and 32GB of RAM. Typical running times for each model selection experiment was approximately 5 minutes without full utilization of the GPU. We ran all experiments (more than 400 combinations) using different methods for multiprocessing in order to use both GPUs at 100% usage. It took less than 6 hours to run all hyper-parameter selection experiments. All other reported results were generated via the Jupyter Notebooks available in the project GitHub.

## 3 Results

### 3.1 Microbial composition is predicted with high accuracy from environmental features

Table 1 shows a quantitative evaluation of our proposed model to satisfy our main goal: to predict the 717 OTU abundances of maize root microbiome starting with some environmental variables.

**Table 1:** Performance of evaluation metrics. In the test set. In Pearson, higher scores are better, because it is a correlation metric. In Bray-Curtis, lower scores are better, as it is a dissimilarity metric.

| Input variables | Default |  | Linear Regression |  | MLP |  | OTU latent space |  | Combined latent space |  |
| --- | --- | --- | --- | --- | --- | --- | --- | --- | --- | --- |
|  | Pearson | Bray-Curtis | Pearson | Bray-Curtis | Pearson | Bray-Curtis | Pearson | Bray-Curtis | Pearson | Bray-Curtis |
| All | 0.5852 | 0.5140 | 0.6629 | 0.4659 | 0.7219 | 0.4169 | **0.7340** | 0.4222 | 0.7229 | **0.4065** |
| Age, T, rain | 0.5852 | 0.5140 | 0.6641 | 0.4638 | 0.6927 | 0.4527 | **0.7348** | 0.4181 | 0.7220 | **0.4072** |
| T and rain | 0.5852 | 0.5140 | 0.5881 | 0.5089 | 0.6323 | 0.5047 | 0.6087 | 0.4686 | **0.6676** | **0.4557** |
| Age and T | 0.5852 | 0.5140 | 0.6622 | 0.4664 | 0.6814 | 0.4591 | **0.7323** | 0.4204 | 0.7189 | **0.4094** |
| Age and rain | 0.5852 | 0.5140 | 0.6628 | 0.4728 | 0.7155 | 0.4211 | **0.7361** | 0.4200 | 0.7048 | **0.4139** |

Given the absence of prior-art or gold standard for this kind of prediction, we compare the accuracy of our models with three baseline models capable of simultaneous regression of multiple variables: a) a default predictor computing the average of all the training samples per each OTU, independently of the mapping features; b) a linear regression model; c) a non-linear model, i.e. a Multi-Layer Perceptron (MLP). In cases b) and c) there is neither latent space, nor an autoencoder; they are alternative approaches to predicting OTU abundances from environmental features.

Table 1 shows that our AE models with latent space (bottom two rows) outperform baseline models (top three rows) in both correlation (e.g., AE models have higher Pearson correlation than non-AE) and community ecology metrics (e.g., the combined model always displays the lowest Bray-Curtis dissimilarity). Notably, OTU latent space models exhibit the best performance.

Among the multiple models tested, on the basis of its performance, we selected the OTU latent space model taking 3 variables (plant age, rain and temperature). This is our reference model for the remainder analysis in the current study. Table 1 shows it is the model with the second best Pearson correlation (with only a slight difference of 0.0013 between first and second-best) and the best Bray-Curtis score among all the OTU latent space models. OTU latent space exhibits the best Pearson correlation for almost all subsets of environmental features (i.e. rows), though the best Bray-Curtis is achieved by the combined latent space models. The reference selected model allows prediction of the microbiome based on changes in both temperature and rain.

### 3.2 Hyperparameters yielding the highest prediction accuracy could be identified

We identified the best combination of hyperparameters resulting in the best model to predict microbial composition, based on a standard validation with a 5-fold CV on the training set.

Our criteria in selecting the optimal model were to both maximize the Pearson correlation, and to minimize Bray-Curtis dissimilarity. The model with the highest Pearson correlation (0.7390 and 0.4204 in Bray-Curtis) corresponded to an architecture with a latent space size of 100. In terms of community ecology, the best model has 0.7384 Pearson and 0.4015 Bray-Curtis, with a latent space of 100. Finally, the best model among those having the smallest latent space size (10) achieves a similar performance to those just described (0.7203 Pearson and 0.4123 Bray-Curtis). As such, in terms of a simpler encoded representation, we selected this as the overall reference model, and used it in the remaining analyses. The detailed parameters and evaluation of the three models described above are in GitHub (experiments 188, 366 and 351 in the ‘autoencoder results’ Jupyter notebook).

Despite our focus on the performance of environmental features in predicting microbial composition, we have also analysed the performance of the reconstruction of the microbial composition from the latent space by our AE. As expected, the performance of the reconstruction alone is better than the performance of the prediction from environmental features (see Figure 2). For example, reaching 0.9174 in Pearson correlation and 0.2065 in Bray-Curtis dissimilarity in 5CV in the reference selected model.

**Figure 2:**
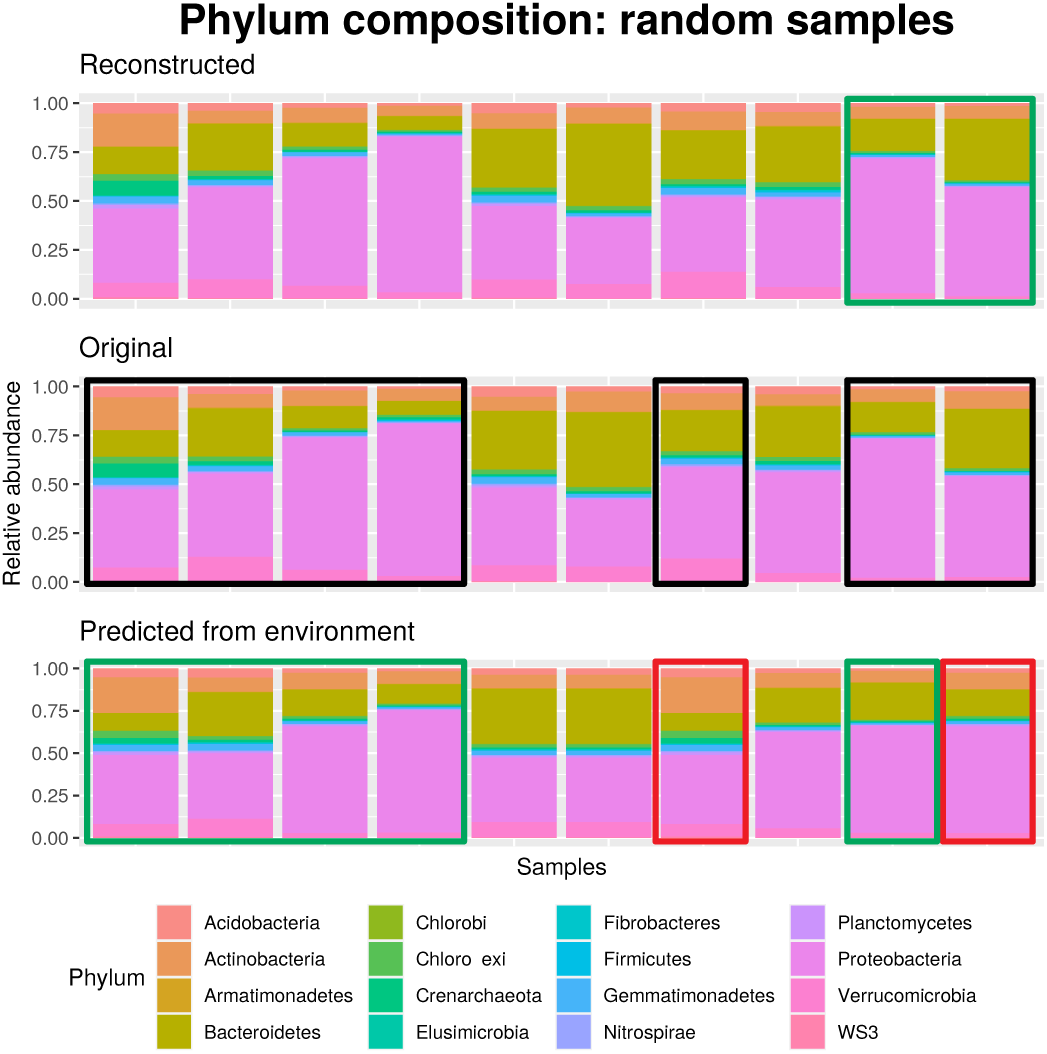
Example of reconstruction and prediction of microbial composition. In the center row is the original microbial composition, allowing it to be compared to both the reconstructed (top) and that predicted from environmental features (bottom). One sample per column. Each Phylum taxonomic category is assigned a different color. Green/Red boxes highlight examples of good/bad sample reconstructions or predictions, and their corresponding original microbial composition is denoted with black boxes.

Regarding the remaining parameters, in normalization, TSS and CLR perform similarly. The larger the latent space the better, though there is only a 0.01 point difference in Pearson correlation between sizes 50 and 100. Smaller autoencoders (i.e. with smaller sizes in the hidden layers) perform better in general; however, we saw the best results with larger autoencoders (more hidden layers, more nodes per layer). We noted little difference between the distinct activation functions. Finally, a batch size of 64 with a learning rate of 0.001 resulted in higher accuracy than a batch size of 128 with a learning rate of 0.01, probably due to the small dataset size used for DL.

The results of the 405 combinations of parameters, with exhaustive metric computations, can be examined in detail in the available ‘autoencoder results’ notebook on GitHub.

### 3.3 Plant age and rain are the most relevant features to predict microbial composition

Table 1 shows, across rows, the performance of different subsets of variables, pointing out that plant age and rain is the combination of features exhibiting the best performance (0.7361 Pearson).

To complement this relevant-features analysis, we trained individual models where the input is only one feature (detailed results available on GitHub). This analysis highlights plant age is the most important feature (0.7330 Pearson), followed by rain (0.6673), temperature (0.6064) and finally inbreds and the maize line (less than 0.58).

Plant age and rain as relevant variables associated to microbial composition are in agreement with the original publication (Walters *et al.*, 2018), although the authors focus in that study was to finding relevant features that enable discovery of heritable OTUs, rather than to predict OTU abundances. The relevance of precipitation to the maize microbiome is also supported by Tan *et al.* (2020), indicating a correlation between that factor and the maize soil bacterial diversity.

### 3.4 *Actinobacteria, Acidobacteria* and *Proteobacteria* were the most accurately predicted taxa

This section explores and interprets the predictive model, providing a qualitative evaluation of the best microbial composition prediction.

Figure 2 shows a comparison of the original (center) vs the reconstructed (top) and environmental-features-predicted microbial composition (bottom) over a demonstrative subset of samples in the test set.

In general, Figure 2 shows a trend towards preserving the pattern of microbial composition (in the figure shown as a similar color distribution pattern) among the three vertical blocks – reconstructed, original and predicted. That tendency is more pronounced in the reconstructed versus the original microbial composition (top and central blocks). For example, the last two samples on the right are very accurately reconstructed (double green box), distinguishing the higher relative abundance of *Bacteroidetes* in the last column. Several other samples, grouped on the left of the figure, show more accurate predictions, as indicated by the green boxes. However, there are samples where the predictions are less accurate (red boxes), for example the last sample on the right, where the abundance of *Bacteroidetes* is particularly badly-predicted. The other red box in Figure 2 highlights another badly-predicted sample, with a higher relative abundance of *Actinobacteria* (orange) in the predicted compared to the original microbiome, and a loss of abundance of *Bacteroidetes, Proteobacteria* (purple) and *Verrucomicrobia* (pink).

In the top 5% predicted OTUs (sorted by ascending Root Relative Squared Error), *Actinobacteria* are the most abundant Phylum group (31.43%), followed by *Acidobacteria* and *Proteobacteria* (22.86% each), *Verrucomicrobia* (8.57%), *Bacteroidetes* and *Planctomycetes* (5.71% each), and *Chloroflexi* (2.86%). Comparatively, the original percentages of OTUs in the test set maize root microbiome show that the most abundant Phyla are *Proteobacteria, Actinobacteria* and *Bacteroidetes* (39.19, 18.83 and 13.39%). Thus, *Bacteroidetes* is a group of taxa under-represented in the best predicted OTUs (5.71%) compared to the reality (13.39%); and, conversely, Acidobacteria is a group of taxa over-represented in the best-predicted OTUs (22.86%) compared to the original OTU data (9.21%).

The ten best-predicted OTUs are (in terms of their Order-level taxonomy): *iii1-15/RB40, MND1, Rhizobiales, 0319-7L14, Ellin6067, Pirellulales, Gaiellales, Actinomycetales, iii1-15* and *Acidimicrobiales*.

### 3.5 Microbiome prediction improves at higher taxonomic levels

Table 2 shows the performance of the prediction of microbial composition when the taxa are aggregated to a higher taxonomic level, reducing the number of OTUs that must be predicted from hundreds to tens. As expected, the higher the taxonomic level (top of the table), the better the performance (i.e. higher Pearson score and lower Bray-Curtis metric). The highest differences are between Phylum and Class, and between Genus and Species.

**Table 2:** Performance at different taxonomic levels. Based on reference model configuration, with the 3 selected input variables.

| Taxonomic level | no. OTUs | OTU latent space |  | Combined latent space |  |
| --- | --- | --- | --- | --- | --- |
|  |  | Pearson | Bray-Curtis | Pearson | Bray-Curtis |
| **Phylum** | 16 | 0.9576 | 0.1591 | 0.9451 | 0.1833 |
| **Class** | 45 | 0.8777 | 0.2514 | 0.8646 | 0.2610 |
| **Order** | 83 | 0.8264 | 0.3007 | 0.7983 | 0.3057 |
| **Family** | 144 | 0.8229 | 0.3239 | 0.7965 | 0.3229 |
| **Genus** | 222 | 0.8133 | 0.3414 | 0.7901 | 0.3408 |
| **Species** | 717 | 0.7348 | 0.4181 | 0.7220 | 0.4072 |

### 3.6 Predicting the microbiome of novel or hypothetical ecosystems

This section describes how our model is able to predict the microbiome of a hypothetical ecosystem, for example, where the environmental features have been defined based on a predicted future environmental state.

Our model has been trained to predict the maize rhizosphere microbiome, using the environmental variables of: temperature, precipitation and plant age.

We designed two challenge scenarios, with the goal of simulating climate change conditions – that is, modifying temperature and humidity. First, we attempt to predict the microbiome in a soil influenced by conditions of higher temperature and lower precipitation (labelled ‘hot and dry’ in Figure 3). Second, we attempt to predict the soil microbiome in “cutoff low” conditions (i.e., low pressure centers aloft) characterized by heavy rains and lower temperatures (labelled ‘cold and wet’ in Figure 3).

**Figure 3:**
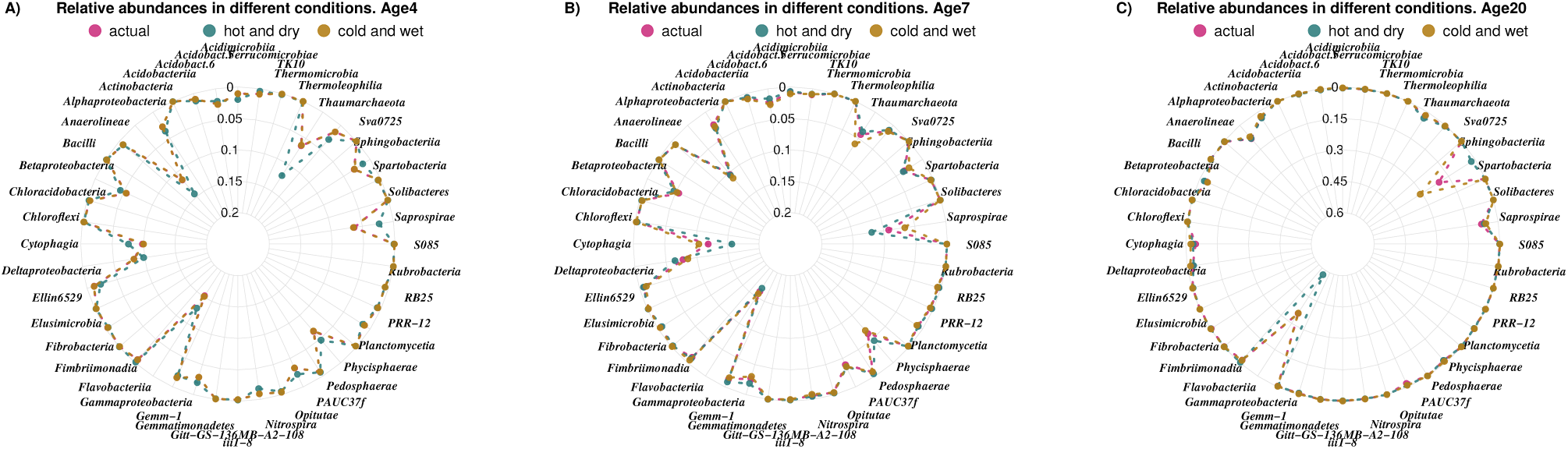
Prediction of microbial composition in different predicted climate change conditions, at distinct plant ages. Outcomes are reported at the Class taxonomic level. Each coloured point and dashed line indicates a sample in a different temperature/precipitation condition. ‘actual’: 59°F and 1.5 inches of rain; ‘hot and dry’: 86°F and 0 inches of rain; ‘cold and rain’: 50°F and 5 inches of rain. Note the difference in the maximum of relative abundance between A/B (0.2) and C (0.6).

Figure 3 compares the relative abundances of the microbial composition (at the Class taxonomic level) in three different conditions: an ‘actual’ situation with intermediate values of temperature and precipitation, and two other scenarios with very high or very low temperature and/or rainfall. Relevant events can be identified when the taxa exhibit divergent relative abundance (i.e., the points in Figure 3 do not overlap), thus corresponding to taxonomic abundance changes predicted to be due to distinct changes in temperature or humidity. Additionally, we analyzed changes at different plant ages.

For example, at an age of 4 weeks (Figure 3A), in ‘hot and dry’ conditions associated to global warming, our model predicts a microbiome with a higher relative abundance of *Thermoleophilia, Alphaproteobacteria* and *Deltaproteobacteria* (green point nearer the center of the radar graph) and a lower relative abundance (i.e. further from the center) of *Saprospirae, Cythophagia, Gammaproteobacteria* and *Pedosphaerae*. This prediction regarding changes under global warming conditions are consistent with previous experimental data about soil microbiomes in arid zones (Liu *et al.*, 2018), where *Thermoleophilia* and *Alphaproteobacteria* appears among the high abundance taxa, and *Gammaproteobacteria* and *Deltaproteobacteria* as low abundance taxa (the latter is the only observation not in agreement with that study). In ‘cold and wet’ conditions, the changes in relative abundances versus actual conditions are less pronounced than in the ‘hot and dry’ condition (the pink and brown points generally overlap or are close). Figure 3B represents the trend of changes in relative abundances of soil microbiome with maize plants at age 7 weeks, under distinct temperature and humidity conditions. Some taxa show a relative abundance gradient over plant age. For example, the relative abundance of *Cythophagia* and *Saprospirae* in ‘hot and dry’ conditions progressively increase into the second month, and eventually become more abundant than the same taxa in the other conditions. In older plants, at 16 or 20 weeks of age (Figure 3C), the relative abundance gradient increases in *Gammaproteobacteria* in ‘hot and dry’ conditions (green), and in *Sphingobacteria* in the ‘cold and wet’ (brown). Additional examples can be seen in the additional ages-result figures available in GitHub.

### 3.7 Transfer learning

In this section we attempt to predict, starting with very few samples, the microbial composition of a maize-associated soil. Our objective is to evaluate the utility of our autoencoder and latent space approach, via transfer learning, in scenarios with a limited number of sequenced microbiome samples (*<*100 being a typical case) and/or a scarce number of measured variables, where *de novo* construction of a predictor would be difficult due to lack of data. In such a scenario, transfer learning from an existing large dataset, onto a smaller, more sparse dataset, would have great utility. Such an approach utilizes the knowledge stored in the latent space (and its decoder).

Our first test of transfer learning is to use the knowledge stored in the deep latent space of our autoencoder, built using a large sample-size, to predict the maize root microbiome from a hypothetical small farm with limited sequencing resources. The hypothetical objective would be that they could be guided towards taking appropriate, contextually-sensitive action (e.g. fertilization, addition of biotics, etc.). To simulate this scenario we took a subset of 100 samples from the Walters *et al.* (2018) dataset (not used previously, neither in training nor test). With that small dataset, we then build a deep latent space predictor from the input features, to be concatenated to the decoder. In this case, therefore, the input variables of the small dataset are the same as those in the model used for the knowledge transfer: temperature, rain and plant age; and the output being the 717 OTUs.

A second test of transfer learning is to assume that the smaller dataset has entirely different environmental features compared to the ones in the primary model. To model this scenario, we took the Maarastawi *et al.* (2018) maize root microbiome dataset. The Maarastawi *et al.* (2018) study did not collect environmental features about weather, but rather about the chemical composition of the soil, so the Maarastawi prediction models use Nitrogen and Carbon concentration, together with pH.

In both transfer learning scenarios, our autoencoder models resulted in better performance than the baseline maize root microbiome predictors, as shown in Table 3. In particular, the model using only the OTU latent space performed best.

**Table 3:** Prediction performance with transfer learning from the primary model to smaller datasets. Walters *et al.* (2018)’s subset: 100 samples with the same input features the primary model for knowledge transfer. Maarastawi *et al.* (2018): 123 samples with environmental features distinct from those in the primary model.

|  | <u>Walters *et al.*’s subset</u> |  | <u>Maarastawi *et al.*</u> |  |
| --- | --- | --- | --- | --- |
|  | Pearson | Bray-Curtis | Pearson | Bray-Curtis |
| **Linear Regression** | 0.5436 | 0.5596 | 0.1588 | 0.6437 |
| **MLP** | 0.7114 | 0.4410 | 0.7230 | 0.3347 |
| **OTU latent sapce** | **0.7544** | **0.4280** | **0.7654** | **0.3233** |
| **Combined latent space** | 0.7266 | 0.4346 | 0.7060 | 0.3728 |

### 3.8 Comparison with similar approaches

Larsen *et al.*, 2012 explored how to predict microbial composition from environmental features; specifically they predicted an oceanic microbiome from environmental variables and interactions (between taxa and with the environment). One important difference when comparing that study with this one is that they reduced the number of taxa at the Species level (hundreds or thousands) by aggregating them at the Order level, resulting in 24 taxa. This resulted in a workable number of output variables, eliminating the need to address the dimensionality reduction problem. In our case, we are able to execute a similar analysis using hundreds of taxa, as shown in Table 2. In contrast to Larsen *et al.*, 2012, we use an Autoencoder architecture. Their intermediate representation was a matrix of interactions between the taxa themselves and between taxa and environmental features, retrieved with a Bayesian Network. The taxa prediction was implemented with a classical Artificial Neural Network, rather than novel DL techniques. Unfortunately, we cannot compare our results quantitatively because neither their data nor their software are available for reproducibility studies.

Ladau *et al.*, 2018 used regression models to establish the relationship between the abundances of the 53 most abundant families in a microbiome, and historical and contemporary climate variables. They concluded that climate change will increase diversity and change the relative abundances of soil bacteria.

From the methodological point of view, Xie *et al.* (2017) designed a similar DL architecture to ours. It is based on an Autoencoder, and is capable of making multiple regressions simultaneously. It is distinct, however, in that it was designed for an entirely different goal: to better understand the mechanisms involved in gene expression regulation. As such, we cannot compare our system – neither quantitatively nor qualitatively – with theirs.

The study most similar to ours is DeepMicro (Oh and Zhang, 2020). Here, we predict a microbiome code from environmental variables and then decode it to obtain the whole microbiome vector using an Autoencoder. DeepMicro also makes use of dimensionality reduction in microbiome data via an Autoencoder architecture; however, the main difference between the approaches is their distinct goals. DeepMicro uses the code to predict human diseases, compared to our approach of predicting the code itself from environmental features, and decoding this code back to a microbiome. These distinct goals make it difficult to do a quantitative comparison of the two approaches. Additional differences include our representation of microbiome composition using only a species-level relative abundance profile, applying different numerical transformations (TSS or CLR), while DeepMicro also tested strain-level marker profiles (0/1 species absence/presence). Further, they use hundreds of samples, while we use thousands in our model training, as is recommended for DL where the objective is to achieve a good estimation of all the hyper-parameter values in the model architecture.

## 4 Discussion

Deep Learning has only rarely been applied to microbiome data due to two main challenges: because the results are difficult to interpret; and the properties of the data are often not favorable for DL. First, interpretability of deep learning models is currently an open research topic of high interest. Some approaches to help interpret deep learning models are beginning to appear, such as the ‘attention approach’, which has been applied to biology (Manica *et al.*, 2019). Second, microbiome data are characterized by high dimensionality (in terms of hundreds or thousands of different microbes) and few samples (usually up to hundreds). Recent large-scale studies are beginning to address these limitations (Sayyari *et al.*, 2019), but they remain as challenges.

First, regarding DL interpretability, we provide a feature relevance analysis. In addition, given that we are solving more than seven hundred regression problems simultaneously (one per each OTU), our performance metrics are aggregated. This would make it more difficult to interpret the results following standard assessments for an unitary prediction problem. To address this, and improve the interpretability of our results, we designed novel dis-aggregated assessments regarding the reliability of an individual microbe prediction (lowest error per OTUs) and the reliability of an individual sample prediction (similarly colored patterns in microbial composition barplots).

Second, our proposal directly addresses the dimensionality problem (i.e. the number of OTUs) by reducing them into the latent space – going from 717 OTUs to 10 values. From there, the problem of predicting microbial composition at low taxonomic levels from a few of environmental features becomes attainable.

Third, another challenge in microbiome data analyses is the low number of samples in an individual study – usually in the low hundreds of samples or less. Our transfer learning proposal contributes a partial solution to that challenge, allowing the building of AI predictive models even with small sample sizes. As we demonstrated, knowledge from the larger Walters *et al.* (2018) maize rhizosphere dataset, with thousands of samples, can be used to transfer predictive knowledge to the similar, but much smaller, Maarastawi *et al.* (2018) study. Moreover, transfer learning can also take advantage of distinct kinds of microbiome metadata, e.g. biotic and abiotic factors – in the case of this study, factors relevant to the soil microbiome such as weather, chemical composition, etc. – combined in the microbiome deep latent space. In such a case, the performance of the prediction with the combined latent space may be expected to improve prediction performance compared to a latent space using just OTUs. This, however, was not observed in the current study, likely because the soil chemical composition factors from the Maarastawi *et al.* (2018) dataset were not measured in the Walters *et al.* (2018) study, thus there was no overlap between the factors studied that could improve prediction performance.

Our approach of creating reduced microbiome representations as a deep latent space could be extended to any other microbiome study type/site (e.g. grapevine, human gut, water, etc.). The requirement is to execute just one large-scale study per study-type or site-type, because DL approaches are only applicable (or recommendable) when many thousands of samples are available. We note that it is possible that rich knowledge in the study meta-data for a particular case may reduce required scale of that initial study.

We note that the requirement to integrate thousands of samples from the same ecosystem/niche in order to build a widely-predictive model also points to a requirement for high-quality meta-data that ensure the datasets are, in fact, comparable and interpretable. This requirement is not always met in practice. These concerns have also been raised by Sakowski *et al.* (2019), who notes that there are few widely accepted quantitative predictive tools in microbiome research, in part due to a lack of standardization, reproducibility, and accessibility of data and methods that would enable undertaking such studies. With the objective of ensuring our work is reproducible, and encouraging better transparency and data publication practices in the community, we have made all of our methods and result files available on GitHub via Jupyter notebooks.

Beyond technical reproducibility of a single study, we also detected more community-wide interoperatibility challenges. For example, between distinct microbiome datasets, due to a lack of standardization in OTU identification and, worse, in the taxonomic labelling/annotation, it is extremely difficult to map results from one study to another. In these cases, starting with the native data, transfer learning is entirely thwarted because there is no overlap in the microbe identifiers whose abundances we want to predict. In this study, we resolved this problem, after a difficult manual intersection between the 717 and 1900 OTUs in the two maize root microbiome datasets, successfully identifying less than half of the expected common OTUs at different taxonomic levels; for example, only 84 of 222 at Genus level (see additional data in section 2.1). Such manual interventions are time-consuming, error prone, and “lossy” (i.e. when a determination of identity cannot be made, the data must be discarded). There is a strong need for the microbiome community to act jointly to establish minimal standards for annotations and for study reporting.

## 5 Conclusions

This manuscript presents, and evaluates, a reproducible deep learning system capable of predicting the microbial composition of the maize rhizosphere starting from only environmental features. It outperforms other approaches in quantitative terms, in particular, providing the advantage of being capable of predicting the relative abundances of hundreds of bacterial species, rather than just a select few.

The computational contributions of our proposed microbiome autoencoder include: a) a novel dimensionality reduction approach to representing microbiome in a reduced latent space; b) the ability to undertake challenging tasks in microbiome data analysis, such as to predict the microbial composition of hundreds of taxa based on a small number of features, rather than the more common (and straightforward) task of predicting a host phenotypic feature from the relative abundance of hundreds of taxa. In addition, this work makes the following contributions to the biological aspect of microbiome analysis: c) the ability to predict microbiome composition based on a limited number of sequenced samples and/or a small number of relevant environmental variables; d) the ability to predict more OTUs than previous approaches, thus enabling predictions at deeper taxonomic levels; and e) the ability to predict hypothetical microbiomes that might arise in novel or predicted/designed ecosystems, for example, under future climate change conditions or after an environmental insult such as a toxic spill.

Our AI system is also explainable. In this study, for example, beyond the biological knowledge arising from an examination of the bacterial species composition, it revealed that the most relevant features to predict this microbial composition are plant age and precipitation. We are also able to identify which taxa are most accurately predicted, in order to be aware of any potential bias of the system.

We then demonstrated that the knowledge in our encoded microbiome can be reused, via transfer learning, to gain insights in distinct but related studies, and by allowing complex analyses to be undertaken using fewer *de novo* sequencing samples. This will be useful in a wide range of cases where data is minimal or unavailable, or where available resources or expertise limit sampling to only a few, easily accessible, environmental features.

This compressed representation opens-up many novel possibilities for microbiome data analysis, particularly with respect to knowledge retrieval and visualization. The condensed representation could be applied to any environment (gut, ocean, urban soil, etc.) where there is a representative set of samples available. The autoencoder structure provides the ability to recover the complete abundance vector from codified samples in the deep latent space, making it possible to perform all analyses using the reduced coded data, and to recover the long vector only when required.

## Acknowledgements

Thanks to the authors of Walters *et al.* (2018) for kindly sharing their datasets with us.

